# Cognition across the lifespan: age, gender, and sociodemographic influences

**DOI:** 10.1101/804765

**Authors:** E. S. Nichols, C. J. Wild, A. M. Owen, A. Soddu

## Abstract

Maintaining cognitive health across the lifespan has been the focus of a multi-billion-dollar industry. In order to guide treatment and interventions, a clear understanding of the way that proficiency in different cognitive domains develops and declines across the lifespan is necessary. Additionally, there are gender differences in a range of other factors, such as anxiety and substance use, that are also known to affect cognition, although the scale of this interaction is unknown. Our objective was to assess differences in cognitive function across the lifespan in men and women in a large, representative sample. Leveraging online cognitive testing, a sample of 18,902 men and women ranging in age from 12-69 matched on socio-demographic factors were studied. Segmented regression was used to model three cognitive domains – short-term memory, verbal abilities, and reasoning. Gender differences in all three domains were minimal; however, after broadening the sample in terms of socio-demographic factors, gender differences appeared. These results suggest that cognition across the lifespan differs for men and women, but is greatly influenced by environmental factors. We discuss these findings within a framework that describes gender differences in cognition as likely guided by a complex interplay between biology and environment.

## Introduction

By 2020, roughly 22% of the world’s population will be over 65, a total of approximately 1.7 billion people (United Nations Department of Economic and Social Affairs, 2019). The consequences of our aging population are many, including an increasing focus on maintaining cognitive health; more so than ever before, individuals are seeking ways to keep their minds sharp. In order to be able to evaluate different tools and treatments for addressing cognitive aging, it is important that we first have a clear understanding of how cognition changes across the lifespan in average, healthy individuals. Additionally, because of the often-cited cognitive differences between women and men (Anderson et al., 2000; Feng et al., 2007; Karapetsas & Vlachos, 1997; Krikorian & Bartok, 1998), we must characterize cognition in each population; if gender differences in cognitive abilities do exist, then men and women may respond differently to cognitive aging interventions.

In healthy individuals, cognitive abilities develop rapidly throughout childhood (Anderson, 2002; Anderson et al., 2001a; Diamond, 2013; Rizeq et al., 2017). By 18, executive function is thought to be mature (Lee et al., 2013), although research suggests that some processes continue to develop in early adulthood (Hartshorne & Germine, 2015). Young adulthood is where most researchers agree that cognitive abilities peak; however there is large variability within this period across different cognitive functions (Anderson, 2002; Hartshorne & Germine, 2015). Mid to late adulthood is then characterized by a slow decline in most cognitive abilities (Diamond, 2013; Salthouse, 2009), and while it can be problematic, this decline is considered part of healthy aging.

Differences in cognitive abilities between men and women are less clear; although several gender disparities in cognitive abilities appear to exist, recent studies have found these differences to be mediated by underlying factors related to gender, such as socio-cultural factors, rather than being inherent to biological factors of sex. For example, Krinzinger and colleagues (2012) found that number processing advantages in boys were mediated by attitudes toward mathematics, and similar results have been found in young adults (Sokolowski et al., 2019). Differences in verbal processing have been less clear, with some suggesting that they are due to variability in instruction and strategy (Scheuringer et al., 2017; Scheuringer & Pletzer, 2017), and others suggesting a hormonal link (Burton et al., 2005; Griksiene & Ruksenas, 2011). Reports of gender differences in age-related cognitive decline are largely thought to be the result of cohort effects (Cornelis et al., 2019; Lipnicki et al., 2017; Wu et al., 2012), although others have found gender-specific links to brain-derived neurotrophic factor (Laing et al., 2012) and brain metabolic activity (Malpetti et al., 2017). Realistically, the truth likely lies somewhere in between, with a multifaceted interaction of biology and environment (Malpetti et al., 2017; Miller & Halpern, 2014).

Finally, there are a number of sociodemographic factors known to affect cognition. For example, it is generally agreed that higher socioeconomic status (SES) predicts better performance on cognitive tasks (Blums et al., 2017; Lubinski, 2009). Additionally, anxiety, depression, and substance abuse also have known detrimental effects on cognition, with higher levels of all three being associated with poorer cognitive outcomes (Crego et al., 2009; Hampshire et al., 2012; Zaremba et al., 2019). Such factors also interact with gender; women tend to experience higher levels of anxiety (McLean et al., 2011) and depression (Parker & Brotchie, 2010), while men experience higher levels of substance abuse (Compton et al., 2007), although women may be more at risk specifically for alcohol abuse (Grant et al., 2017, but see Bratberg et al., 2016). Thus, there is a complex interaction of age, gender, and other sociodemographic variables that must be considered when studying cognitive abilities across the lifespan.

The internet provides a unique opportunity for examining cognition across the lifespan in the general population on a huge scale, allowing data to be sampled from participants from a broad range of SES, geographical, and educational backgrounds. Leveraging the power of the internet provides us with a cross-sectional snapshot of both demographics and cognition from a larger and more diverse sample than would be possible to collect in the laboratory.

The first goal of the present study was to characterize cognitive abilities across the lifespan, ranging from adolescence to late adulthood. Specifically, we sought to address whether differences exist between cognitive domains; do different cognitive domains show the same pattern, or are they at their peak at different ages? Do they show the same rate of decline, or do some remain resilient to aging more so than others? The second goal was to examine whether age effects differed between genders, and what factors may influence these differences. Specifically, do gender differences exist in some cognitive domains and not others? Do men and women attain their highest scores at the same age, and do they decline at the same rate? Further, we explored the demographic and social factors that affect the genders differently, and whether controlling for these differences affects the observed pattern of cognitive abilities across the lifespan. Taking into account studies of the effects of mental health and sociodemographic variables on cognition, we predicted that matching groups on these factors would eliminate gender differences in cognitive abilities. However, based on smaller studies using more limited time windows, we predicted that when not controlling for these factors, gender differences would manifest with men outperforming women in memory and reasoning, but with women outperforming men in verbal abilities, and that the pattern of these abilities would show an increase up to early adulthood, and a slow decline into mid and late adulthood.

## Materials and Methods

### Participants

All data for this study were collected with the CBS (www.cambridgebrainsciences.com) online platform, which has previously been used for other large-scale studies of cognition (Nichols et al., 2020; Wild et al., 2018). From a database of 65,994 participants, two tightly matched samples of men and women were created, with 9,451 participants in each. A summary of the sample’s demographics is included in Table 1. All participants gave informed consent, and ethics approval was obtained through the local Research Ethics Committee (2010.62).

**Table 1.**
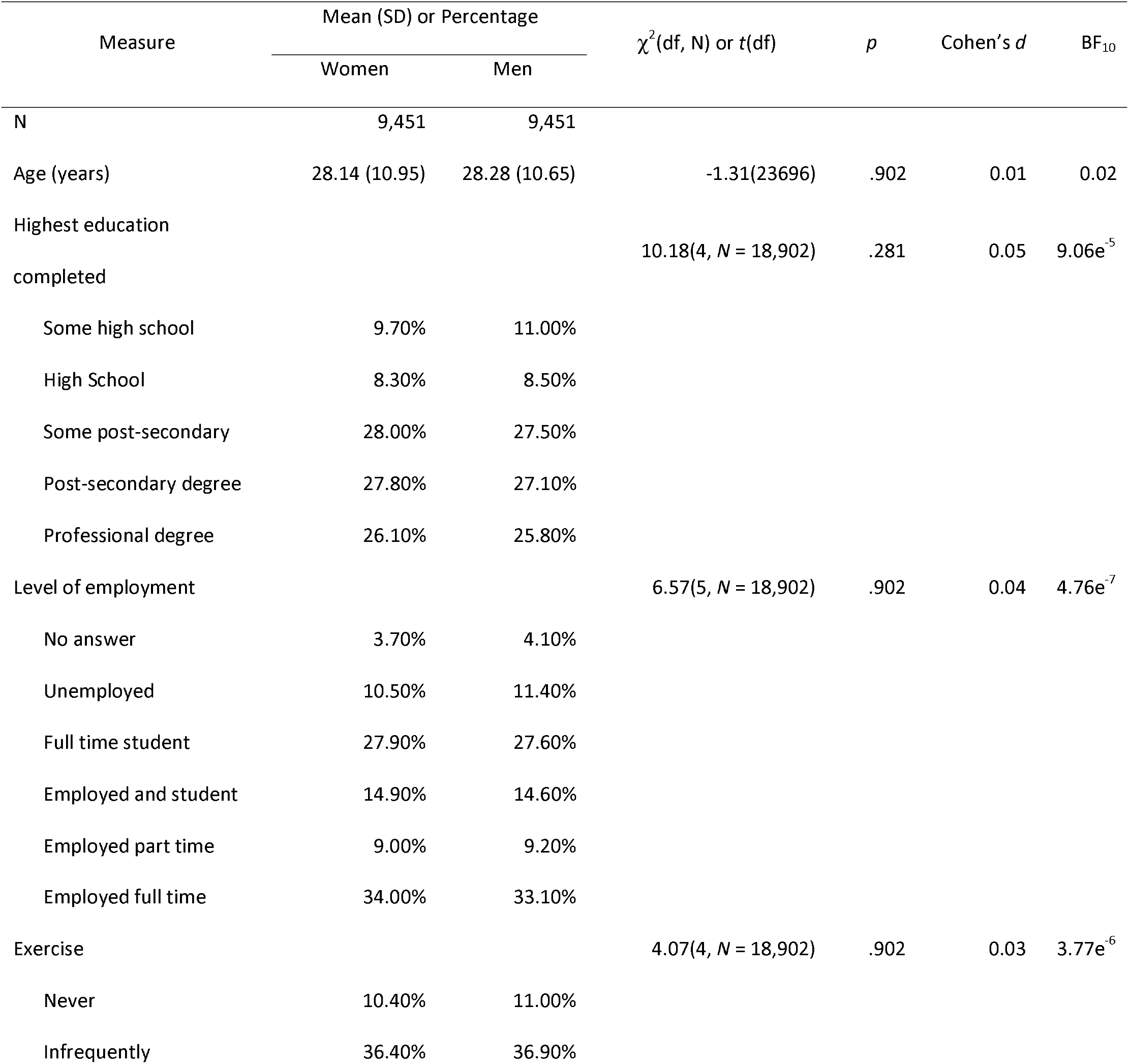
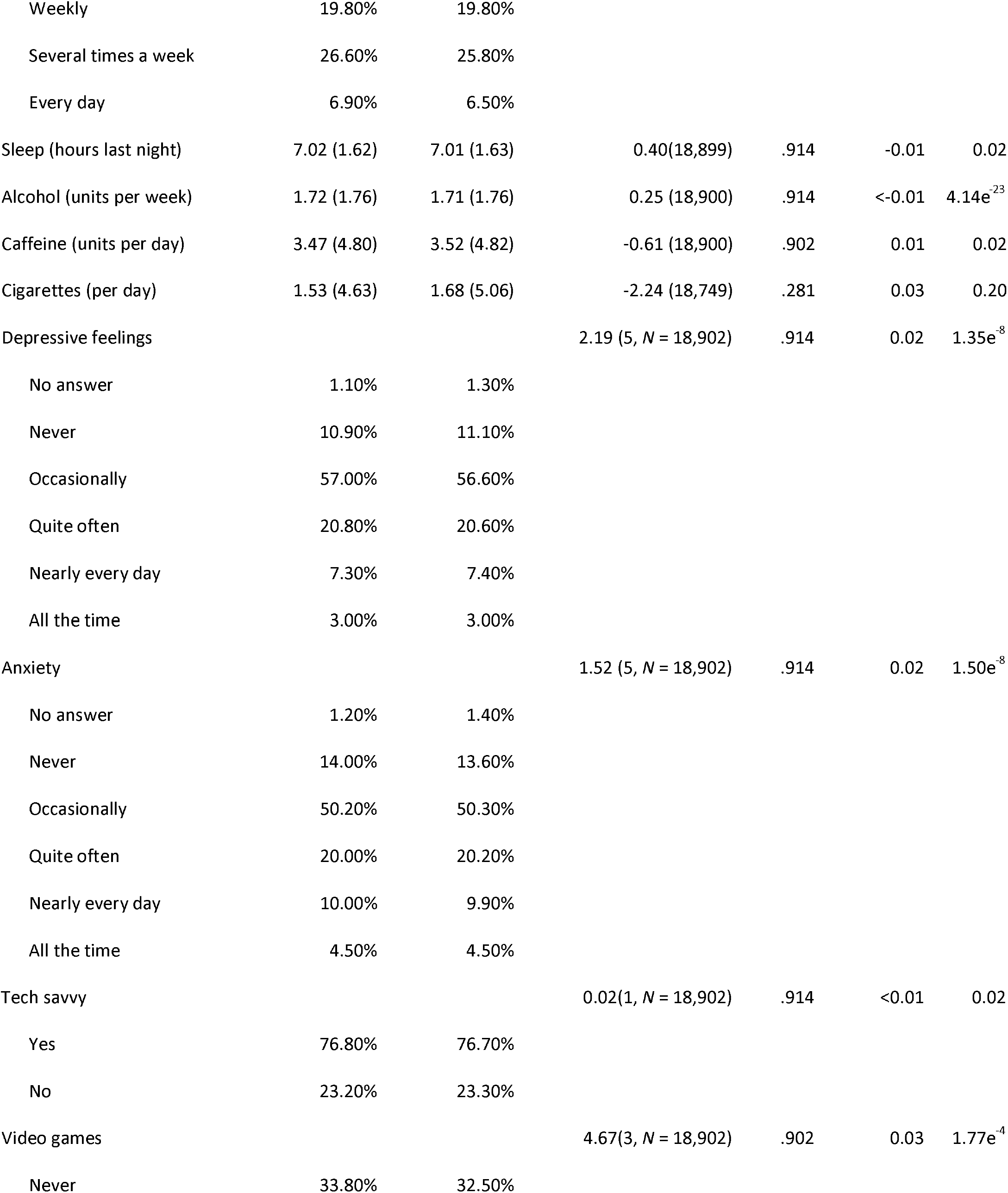
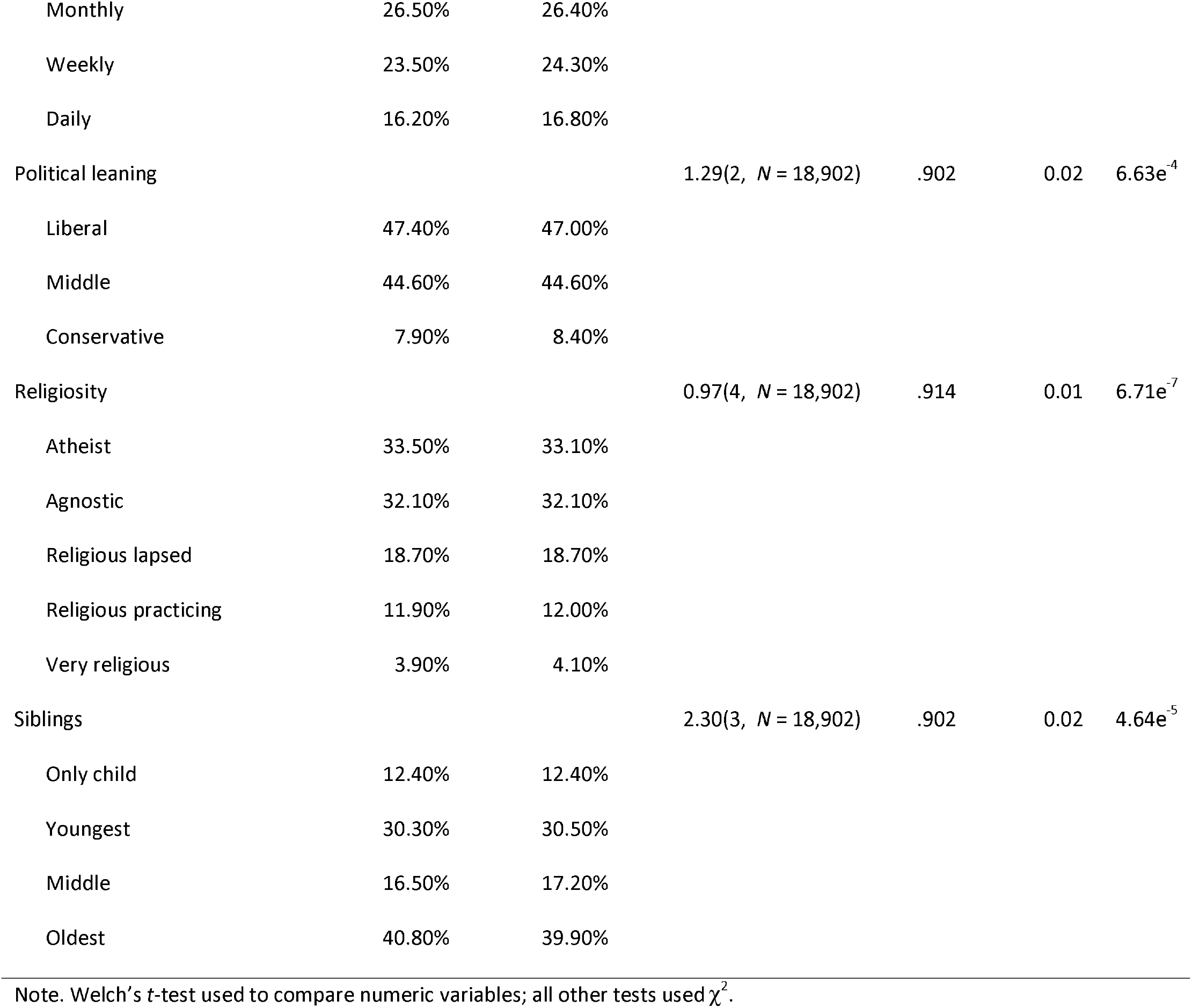
Comparison of demographic variables across women and men

### Materials

#### Sociodemographic, lifestyle, psychological, and sleep questionnaire

The sociodemographic, lifestyle, psychological, and sleep questionnaire included questions about the individual’s age and gender, lifestyle such as exercise, substance use, and sleep, mental health such as depressive symptoms and anxiety, and other information such as education, employment, and level of technical savviness. When these data were collected, gender was presented as a binary response (male/female), therefore we do not have information on non-binary individuals. Data included in the present study are listed in Table 1. The questions used in the present study are included in the Supplementary Material.

#### Cognitive battery

Prior to filling in the questionnaire, participants completed the 12 tests in the CBS battery. Test order was fixed across participants. Detailed descriptions of the tests can be found in the Supplementary Material, but in brief they are: (1) ‘Monkey Ladder’ (visuospatial working memory); (2) ‘Grammatical Reasoning’ (verbal reasoning); (3) ‘Double Trouble’ (a modified Stroop task); (4) ‘Odd One Out’ (deductive reasoning); (5) ‘Spatial Span’ (short-term memory); (6) ‘Rotations’ (mental rotation); (7) ‘Feature Match’ (feature-based attention and concentration); (8) ‘Digit Span’ (verbal working memory); (9) ‘Spatial Planning’ (planning and executive function); (10) ‘Paired Associates’ (shape-location associative memory); (11) ‘Interlocking Polygons’ (visuospatial processing); and (12) ‘Token Search’ (working memory and strategy).

### Factor analysis

The 12 tests were used to create three “composite” scores reflecting performance based on a previous factor analysis described in Hampshire et al. (2012). The three composite scores, labeled as short-term memory, reasoning, and verbal abilities, were calculated as follows. First, the individual test scores were normalized *(M = 0.0, SD = 1.0).* Then, the three cognitive domain scores were calculated using the formula *Y* = *X*(*Ar*^+^)^T^, where *Y* is the N × 3 matrix of domain scores, *X* is the N × 12 matrix of test z-scores, and *Ar* is the 12 × 3 matrix of varimax-rotated principal component weights from Hampshire et al. All 12 tests contributed to each domain score, as determined by their component weights.

### Statistical analyses

Data were analyzed in R (version 3.5.2, R Core Team, 2018) and RStudio (version 1.1.463). Specific packages included: ‘Segmented’ (Muggeo, 2008) for computing regressions with breakpoints, ‘Matchit’ (Ho et al., 2011) for matching samples on demographic variables, ‘parallel’ for parallel computing, and ‘boot’ (Canty & Ripley, 2019) for calculating confidence intervals. Figures were produced using ‘ggplot2’ (Wickham, 2016). Two groups of 9,451 men and 9,451 women were created, matched on with the nearest neighbour matching method for all variables listed in Table 1.

To examine the differences in demographic variables between genders, three different tests were used: Welch’s *t*-tests for continuous variables, Wilcoxon Rank Sum tests for ordinal variables, and chi-square tests for categorical variables. P-values were corrected for multiple comparisons using a false discovery rate and were considered significant at *p* < .01. Effect size was calculated using the appropriate measures for each test: Cohen’s *d* for *t*-tests, rfor Wilcoxon Rank Sum tests, and Cramer’s Vfor chi-square tests. Measures of skew and kurtosis indicated that domain scores were normally distributed, and histograms are shown in Figure 1.

**Figure 1.**
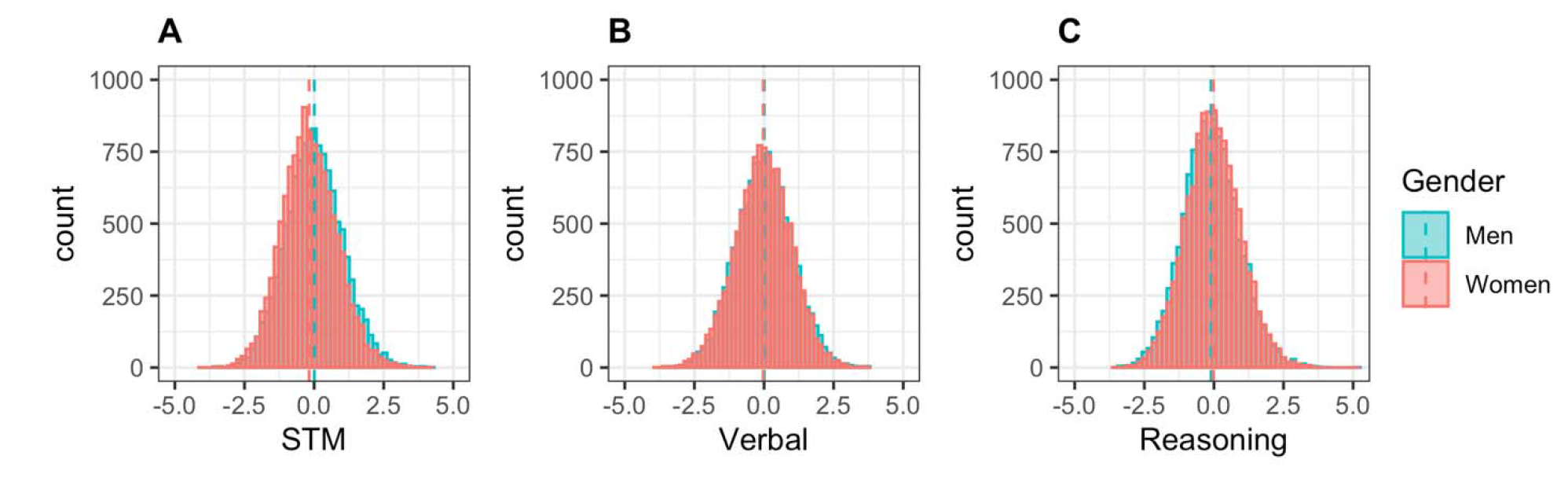
Histograms of domain scores by gender. Dashed lines indicate mean.

Segmented linear regression models were constructed to predict each of the 3 domain scores from participants’ reported age and were estimated using maximum likelihood estimation. Segmented regression was used to fit a model in which there is a change in the linear relationship – such as a “peak” that indicates a transition from increasing to decreasing performance across different ages – without imposing a pre-determined shape (e.g., quadratic or cubic) through adding one or more piecewise linear relationships (Muggeo, 2003, 2008). The value of the independent variable (i.e., age) at which this change occurs is referred to as a breakpoint. The relationship between cognitive performance and age was modeled separately for each gender.

The segmented regression technique used here requires that the number of breakpoints, and (optionally) initial estimates of their locations, are provided. To determine the number of these points in each score, we fit each segmented regression model multiple times with one or more breakpoints and selected the model with the lowest Bayesian Information Criterion (BIC) (Muggeo, 2008; Tiwari et al., 2005). The number of breakpoints was estimated separately for each domain score and gender. The algorithm converged on consistent breakpoint locations regardless of whether initial estimates were provided (from visual inspection of local regression curves, shown in Figure S1), or not. To confirm that a model with one or more breakpoints predicted the data better than a linear model, the Davies’ test (Davies, 2002) was used to determine whether there was a statistically significant change in slope. The estimated breakpoint location was taken as the age that was associated with peak performance in all regression models except for two cases. First, in men’s verbal scores, in which there were two breakpoints and the breakpoint with the highest score was used as the age at which performance peaked. Second, in women’s reasoning scores, in which the highest score was at the lower boundary of our age range. Slopes of the increasing and decreasing segments, as well as the middle segment for men’s verbal scores, were obtained using the ‘slope’ function of the ‘segmented’ package, and 95% confidence intervals (CIs) were calculated for peak age, score at peak age, and all slopes.

Differences in these parameters between men and women were analyzed by bootstrapping with 10,000 replications the difference of the estimated parameter values from models that were separately estimated for men and women. To determine whether these values differed significantly between genders, the lower and upper 2.5% quantiles of the bootstrapped difference values were produced; if these bounds included zero, then it could be interpreted as no significant difference between the genders.

In segmented models where multiple breakpoints were deemed a better solution than a single point as determined using BIC, the increasing or decreasing portion of the curve (i.e., the data to the left or right of the “peak”) was characterized by two increasing or decreasing linear segments with different slopes (as can be seen in Figure 2C, women’s reasoning scores). In order to compare slopes between the genders in these cases, bootstrapping was conducted by fitting the segmented model, then calculating the average slope to the left (in the case of men’s verbal scores) or right (in the case of women’s reasoning scores) of the peak. The rest of the bootstrapping parameters were kept the same as described above.

**Figure 2.**
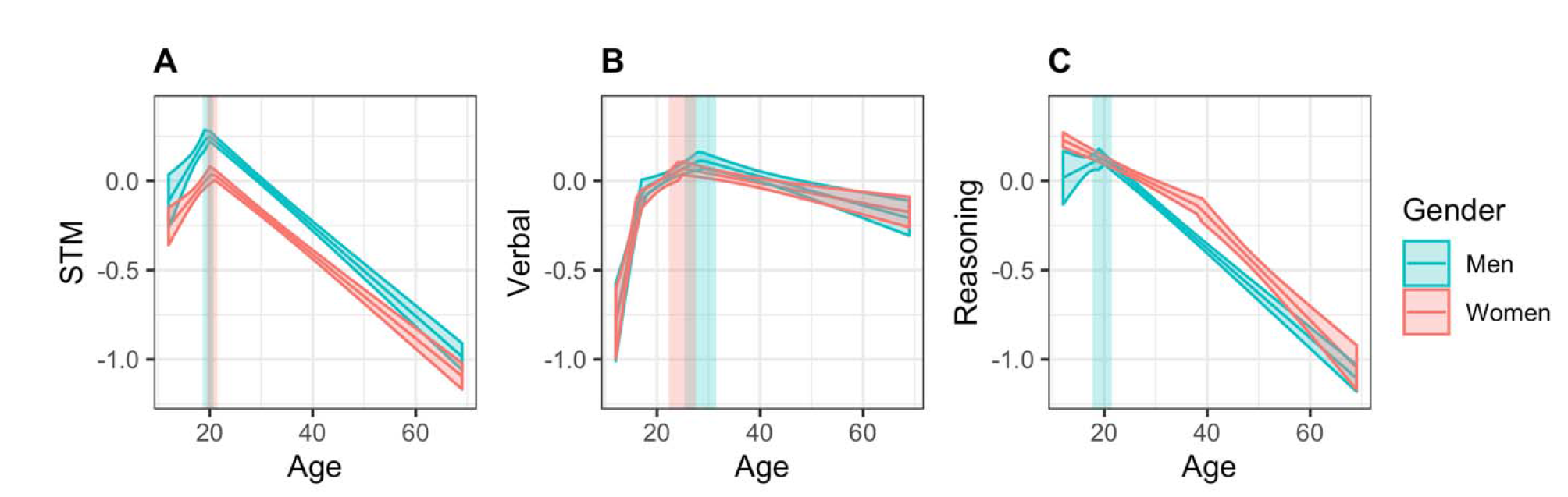
Regression lines for STM, Verbal, and Reasoning scores across the lifespan, ranging from 12 to 69 years of age. 95% simultaneous confidence bands are shown in translucent colour around the line, and 95% confidence intervals for peak age are shown in translucent rectangles.

### Secondary analyses

Although matching groups on sociodemographic measures allows us to more accurately determine what the influence of gender alone is on cognitive performance, men and women do realistically differ on measures such as anxiety and sleep, and such factors are known to affect cognition. Thus, a second set of analyses were run on the full database (after cleaning of missing data and outliers, described below), to determine what differences may exist in a sample that is reflective of the sociodemographic variance we see in the population.

Only data from the participants who completed all questionnaire items and all 12 tests were included in analysis. 65,994 participants met these requirements. Test scores were then filtered for outliers in two passes: scores greater than six standard deviations were assumed to be technical errors and were first removed. Then, scores greater than four standard deviations from the recalculated mean were identified, assumed to be performance outliers, and removed. Finally, individuals younger than 12 and older than 69 were removed because of low numbers outside of this age range. 45,779 participants were included in the final analysis.

Descriptive information for these two new samples is summarized in Table S1. Scores are plotted against age in Figure S2, and histograms of domain scores are shown in Figure S3. Local regression curves are shown in Figure S4. The same set of analyses were performed as outlined in the section above, however because the total sample of men was larger than women, a random sample of 13,444 men were selected upon each bootstrap iteration in order to match the female sample size.

## Results

### Cognitive domain scores

#### Short-term memory

Results are reported in Table 2. A model with one breakpoint was found to best estimate women’s memory scores. The highest point in women’s STM scores occurred at age 20.42 [95% CI = 19.36, 21.48], with a score of 0.046 [95% CI = −0.009, 0.101]. The slopes of the segments to the left and right of the breakpoint were 0.036 [95% CI = 0.019, 0.053] and −0.023 [95% CI = −0.025, −0.022], respectively, indicating that age was a significant predictor of STM performance in these age ranges; specifically, increasing age was associated with increasing scores up to the age of 20 years, after which it was associated with decreasing performance. Davies’ test for a change in slope was significant (p < .001), indicating that the linear relationship changed at the breakpoint, as can be seen in Figure 2A.

**Table 2:** Segmented regression parameter estimates for age, from regression models estimated for each composite score

| Score | Gender | Term | Coef | SE | *t* | *p* |
| --- | --- | --- | --- | --- | --- | --- |
| STM | Women | Age | 0.04 | 0.01 | 4.10 | < .001 |
|  |  | ΔAge | -0.06 |  |  |  |
|  | Men | Age | 0.05 | 0.01 | 3.61 | < .001 |
|  |  | ΔAge | -0.07 |  |  |  |
| Verbal | Women | Age | 0.15 | 0.01 | 7.58 | < .001 |
|  |  | ΔAge1 | -0.13 |  |  |  |
|  |  | ΔAge2 | -0.03 |  |  |  |
|  | Men | Age | 0.15 | 0.03 | 5.36 | < .001 |
|  |  | ΔAge1 | -0.13 |  |  |  |
|  |  | ΔAge2 | -0.02 |  |  |  |
| Reasoning | Women | Age | -0.01 | 0.001 | -8.83 | < .001 |
|  |  | ΔAge | -0.02 |  |  |  |
|  | Men | Age | 0.01 | 0.01 | 1.10 | .272 |
|  |  | ΔAge | -0.04 |  |  |  |

Note: *p*-values for change in slope measured by Davies' test; ΔAge refers to change in age parameter after a breakpoint

Men’s memory scores were also best estimated by a segmented model with one breakpoint. The highest point in men’s STM score occurred at age 19.65 [95% CI = 18.61, 21.48], with a score of 0.259 [95% CI = 0.187, 0.330]. Slope of the increasing segment was 0.049 [95% CI = 0.022, 0.075], and slope of the decreasing segment was −0.025 [95% CI = −0.027, −0.023], showing a significant effect of age on STM score in men. The change in slope was significant, as measured by the Davies’ test (p < .001). As can be seen in Table 3, there was no significant difference in the age at which women and men peaked in STM performance. However, men reached a significantly higher overall score than women at their peak ages, a difference of 0.21 standard deviations. When comparing how STM scores increased leading up to peak age and how quickly they declined afterward, women and men did not differ significantly.

**Table 3:** Comparisons between genders matched on socio-demographic variables

| Score | Measure | Women [95% CI] |  | Men [95% CI] |  | Difference [95% CI] |  |
| --- | --- | --- | --- | --- | --- | --- | --- |
| STM | Peak age | 20.42 | [19.36, 21.48] | 19.65 | [18.61, 20.69] | 0.76 | [-2.09, 4.32] |
|  | Peak score | 0.046 | [-0.009, 0.101] | 0.259 | [0.187, 0.330] | -0.213 | [-2.63, -0.159] |
|  | Increase | 0.036 | [0.019, 0.053] | 0.049 | [0.022, 0.075] | -0.013 | [-0.132, 0.028] |
|  | Decrease | -0.023 | [-0.025, -0.022] | -0.025 | [-0.027, -0.023] | 0.002 | [-0.001, 0.005] |
| Verbal | Peak age | 24.89 | [22.26, 27.52] | 28.42 | [25.33, 31.52] | -3.53 | [-20.49, 6.10] |
|  | Peak score | 0.071 | [0.033, 0.108] | 0.104 | [0.050, 0.158] | -0.033 | [-0.091, 0.019] |
|  | Increase | 0.035 | [0.016, 0.048]<sup>a</sup> | 0.022 | [0.006, 0.045]<sup>a</sup> | 0.013 | [-0.012, 0.036] |
|  | Decrease | -0.006 | [-0.008, -0.003] | -0.008 | [-0.011, -0.005] | 0.002 | [-0.003, 0.014] |
| Reasoning | Peak age | 12 |  | 19.62 | [17.70, 21.54] | -7.62 | [-12.82, -2.23] |
|  | Peak score | 0.223 | [0.187, 0.271] | 0.131 | [0.060, 0.201] | 0.092 | [-0.047, 0.151] |
|  | Increase | — |  | 0.015 | [-0.012, 0.041] | — |  |
|  | Decrease | -0.020 | [-0.021, -0.018]<sup>a</sup> | -0.025 | [-0.027, -0.023] | 0.005 | [0.003, 0.008] |

<sup>a</sup> Combined slope across two segments is reported.

#### Verbal abilities

Results of segmented regression of verbal scores are also summarized in Table 2. A model with two breakpoints was found to best estimate women’s verbal scores. Women first had a breakpoint at age 16.49, at which point the rate at which scores were increasing, slowed (Figure 2B). The highest point in women’s verbal scores occurred at age 24.89 [95% CI = 22.26, 27.52] with a score of 0.071 [95% CI = 0.033, 0.108]. Slope of the initial increasing segment was 0.153 [95% CI = 0.093, 0.214], the slope of the second increasing segment was 0.022 [95% CI = 0.009, 0.035] and slope of the decreasing segment was −0.006 [95% CI = −0.008, −0.003], showing a significant relationship between age and verbal abilities. Davies’ test for a change in slope was significant (p < .001), indicating that the linear relationship changed at the breakpoint.

Men’s verbal scores were best estimated by a segmented model with two breakpoints. As can be seen in Figure 2B, men first had a breakpoint at age 17.16, at which point the rate at which scores were increasing, slowed. The highest point in men’s verbal score occurred at age 28.42 [95% CI = 25.33, 31.52], with a score of 0.104 [95% CI = 0.050, 0.158]. Slope of the initial increasing segment was 0.146 [95% CI = 0.094, 0.198], the slope of the second increasing segment was 0.015 [95% CI = 0.006, 0.023] and slope of the decreasing segment was −0.008 [95% CI = −0.011, −0.005], indicating a significant relationship between age and verbal abilities in all three sections. The change in slope was significant, as measured by the Davies’ test (p < .001).

As summarized in Table 3, there were no significant differences in the age at which women and men’s scores reached a maximum in verbal abilities, scores at peak age, nor in the slopes of the increase and decrease in scores surrounding peak age

#### Reasoning

A model with one breakpoint was again found to best estimate women’s reasoning scores. However, this breakpoint occurred at age 38.12 years, and indicated a transition from a gradual to steeper decline: scores declined with a slope of −0.014 [95% CI = −0.017, −0.011] from age 12 to age 38.12, at which point the negative slope increased to −0.029 [95% CI = −0.035, – 0.024]. Davies’ test for a change in slope was significant (p < .001), indicating that the linear relationship changed. As can be seen in Figure 2C, the highest predicted scores for women occurred at age 12 with a score of 0.223 [95% CI = 0.187, 0.271]. However, because this is the cut-off age of our sample, it is not possible to determine whether this is indeed a true peak, or if scores are higher at earlier ages.

Men’s reasoning scores were best estimated by a segmented model with one breakpoint. The breakpoint in men’s reasoning score occurred at age 19.62 (95% CI = 17.70, 21.54), with a score of 0.131 [95% CI = 0.060, 0.201]. The change in slope was significant, as measured by the Davies’ test (p < .001), however the slope of the initial segment was 0.015 [95% CI = −0.012, 0.041], and slope of the decreasing segment was −0.025 [95% CI = −0.027, – 0.023], indicating that only the second segment showed a significant effect of age. Similar to women, this suggests that we did not capture a developmental increase in reasoning abilities within the current sample, and it is possible that the true peak occurs earlier than age 12.

Because we do not have a reliable measure of peak age in either gender, we compared between genders the age at which reasoning scores began to decline. Women began to decline significantly earlier than men, however reasoning scores at that age did not differ between genders (Table 3). Because women did not show an increase in reasoning scores within our age range, we could not compare men and women on this measure. However, when comparing how scores declined after peak age, men declined significantly faster than women.

### Unmatched samples

Women and men differed on several demographic factors, but not for age, education, exercise, and number of siblings. While all significant p-values were ≤ .003, the largest effect sizes were seen in hours of sleep (Cohen’s *d =* 0.10), units of caffeine per day (Cohen’s *d = −* 0.19), anxiety level (Wilcoxon’s *r =* 0.15), and technical savviness (Cramer’s *V=* 0.24).

#### Short-term memory

Results of the segmented regression for STM scores of both genders in the socio-demographically unmatched sample are reported in Table 4. Both women and men showed a significant change in slope as measured by the Davies’ test (p < .001 for both genders). As can be seen in Table 5 and Figure 3A, no significant differences were found in the age at which women and men reached the highest point in STM, nor in the slopes of the increase and decrease in scores surrounding peak age. However, men reached a higher overall score than women at their peak ages by a standard deviation of 0.28.

**Figure 3.**
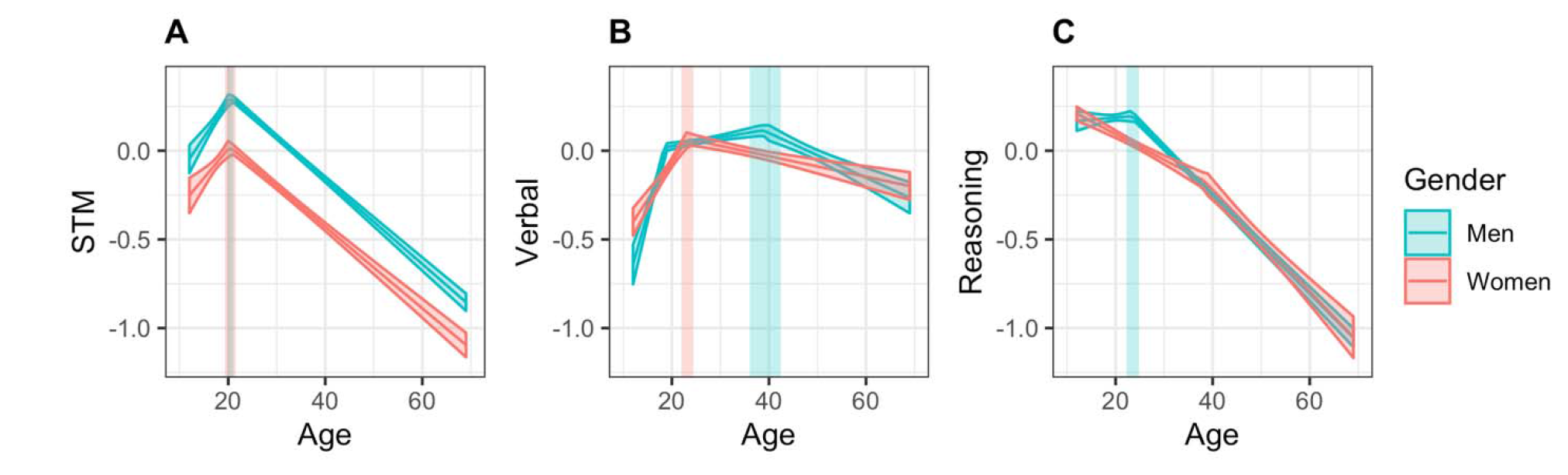
Regression lines for STM, Verbal, and Reasoning scores across the lifespan, ranging from 12 to 69 years of age, in the socio-demographically unmatched sample. 95% simultaneous confidence bands are shown in translucent colour around the line, and 95% confidence intervals for peak age are shown in translucent rectangles.

**Table 4:** Segmented regression parameter estimates for age, from regression models estimated for each composite score, for models estimated with N = 45,779

| Score | Gender | Term | Coef | SE | *t* | *p* |
| --- | --- | --- | --- | --- | --- | --- |
| STM | Women | Age | 0.03 | 0.01 | 3.83 | < .001 |
|  |  | ΔAge | -0.05 |  |  |  |
|  | Men | Age | 0.04 | 0.01 | 6.48 | < .001 |
|  |  | ΔAge | -0.07 |  |  |  |
| Verbal | Women | Age | 0.04 | 0.01 | 8.16 | < .001 |
|  |  | ΔAge | -0.05 |  |  |  |
|  | Men | Age | 0.10 | 0.01 | 8.44 | < .001 |
|  |  | ΔAge1 | -0.09 |  |  |  |
|  |  | ΔAge2 | -0.02 |  |  |  |
| Reasoning | Women | Age | -0.01 | 0.001 | -9.62 | < .001 |
|  |  | ΔAge | -0.01 |  |  |  |
|  | Men | Age | 0.003 | 0.004 | 0.73 | .468 |
|  |  | ΔAge | -0.03 |  |  |  |

Note: *p*-values for change in slope measured by Davies' test; ΔAge refers to change in age parameter after the breakpoint

**Table 5:** Comparisons between genders on key measures of cognitive performance over the lifetime, for models estimated with N = 45,779

| Score | Measure | Women [95% CI] |  | Men [95% CI] |  | Difference [95% CI] |  |
| --- | --- | --- | --- | --- | --- | --- | --- |
| STM | Peak age | 20.47 | [19.39, 21.55] | 20.48 | [19.85, 21.12] | -0.01 | [-4.70, 3.44] |
|  | Peak score | 0.021 | [-0.007, 0.049] | 0.304 | [0.286, 0.323] | -0.283 | [-0.331, -0.219] |
|  | Increasing slope | 0.032 | [0.015, 0.048] | 0.042 | [0.029, 0.054] | 0.010 | [-0.071, 0.036] |
|  | Decreasing slope | -0.023 | [-0.025, -0.021] | -0.024 | [-0.025, -0.023] | 0.001 | [-0.002, 0.005] |
| Verbal | Peak age | 23.21 | [22.00, 24.42] | 39.20 | [35.99, 42.42] | -15.99 | [-26.36, -3.86] |
|  | Peak score | 0.067 | [0.033, 0.101] | 0.116 | [0.074, 0.157] | -0.049 | [-0.145, -0.002] |
|  | Increasing slope | 0.042 | [0.032, 0.052] | 0.014 | [0.007, 0.027]<sup>a</sup> | 0.028 | [0.012, 0.176] |
|  | Decreasing slope | -0.006 | [-0.008, -0.004] | -0.013 | [-0.017, -0.009] | 0.007 | [-0.001, 0.019] |
| Reasoning | Peak age | 12 |  | 23.51 | [22.25, 24.78] | -11.51 | [-16.96, -4.22] |
|  | Peak score | 0.208 | [0.168, 0.249] | 0.196 | [0.163, 0.228] | 0.012 | [-0.136, 0.046] |
|  | Increasing slope | — |  | 0.003 | [-0.004, 0.010] | — |  |
|  | Decreasing slope | -0.019 | [-0.021, -0.018]<sup>a</sup> | -0.027 | [-0.029, -0.026] | 0.008 | [0.004, 0.012] |

Note: Values are missing for women's reasoning increasing slope as both segments were negative
<sup>a</sup> Combined slope across two segments is reported. Slopes of the individual segments are reported in-text.

#### Verbal abilities

Both women and men showed a significant change in slope as measured by the Davies’ test (p < .001 in all tests). A model with a single breakpoint best estimated women’s scores, while men’s scores were still estimated best by a model with two breakpoints. As summarized in Table 5, men reached the highest point in verbal abilities at a significantly later age than women. Men also had significantly higher scores at peak age, with a difference of 0.05 standard deviations. When comparing how scores increased up to peak age, women’s scores improved at a faster rate than men’s, however there was no difference when comparing the rate of decline from peak age to age 69.

#### Reasoning

Reasoning scores in our sample of women began to decrease at a significantly earlier age than men, however scores at that age did not differ between genders. While we did not capture an increase in reasoning abilities in either gender in our sample, reasoning scores decreased significantly faster in men than women (Table 5).

## Discussion

After creating three cognitive domain scores from the 12 cognitive tests based on their underlying factor structure, we replicated previous findings that not all cognitive domains develop and decline in the same way. Specifically, STM increased rapidly from age 12 to the early 20s, at which point it decreased at a steady rate until age 69, the upper limit of our sample’s age range. Verbal abilities also peaked in early adulthood, while reasoning did not show a clear peak in scores, instead being characterized by either a decline from age 12, or a plateau followed by a decline. These results were consistent with previous studies showing that cognition is not a unitary concept, and different cognitive abilities have separable developmental trajectories (Hartshorne & Germine, 2015; Salthouse, 2009). However, they extend the results of those studies in several important ways:

Interpreting gender differences in cognitive data is complicated by the differences in socio-demographic factors. Several factors that were matched across groups, such as sleep and anxiety, have known effects on cognitive function (Wild et al., 2018), making it difficult to determine what is driving the observed gender differences in samples unmatched on these variables. Additionally, because these socio-demographic factors are gender-dependent, it is not possible to include them in the model due to issues with multicollinearity. By matching men and women on these factors, however, we were able to limit their effect on the data as much as possible, and this greatly reduced or eliminated the differences in cognitive performance and aging. Of course, there are numerous factors that we did not control for, such as reproductive health and occupation, and it is impossible to truly capture all of them. Additionally, there are socio-demographic differences that may have biological underpinnings. For example, depression is more prevalent in women, perhaps due to the presence of sex-specific forms such as premenstrual dysphoric disorder (Albert, 2015). It is therefore difficult to disentangle the environment from biological sex differences, however accounting for these differences, regardless of their origin, is necessary for describing gender differences in cognition alone.

While these results are presumed to be reflective of the cognitive performance in a tightly controlled sample, when examining the progression of STM, verbal abilities, and reasoning in men and women in the broader database, all three cognitive domains showed unique differences. Although men and women’s scores reached peak STM performance at the same age, men reached a slightly higher score than women. In verbal abilities, women peaked faster and earlier, but men again reached higher scores. While women’s reasoning began to decline earlier than men’s, men declined at a faster rate. These results extend what is known from previous gender research. For example, there is evidence that men lose grey matter volume more rapidly with age than women, especially in fronto-temporal regions (Kryspin-Exner et al., 2011; A. K. H. Miller et al., 1980; Sowell et al., 2007); this in turn may lead to faster decline in cognitive function, fitting the pattern observed here in the reasoning domain. In contrast, women are thought to have better verbal processing than men; however we see the opposite here, with men reaching a higher peak score than women. One possible explanation for this discrepancy could be the age at which verbal abilities are tested. Burton and colleagues (19) tested a sample of university students, which is common in Psychology research. Looking at the pattern of verbal abilities in men and women in the current unmatched sample, women seem to outperform men at age 23, which, if we were to only examine individuals around this age, may lead to the erroneous conclusion that women have superior verbal abilities. Similarly, men are frequently reported to be better at mental rotation than women (Burton et al., 2005), a test included in our reasoning domain. Here, we found that peak reasoning scores did not differ between genders, but women declined much earlier than men. Again, comparing genders within a limited age range would have led to the erroneous conclusion that men outperform women in this domain, when in reality it is a difference in trajectory of reasoning abilities. The present results underline the need to take the progression of cognitive abilities across the lifespan into account when studying gender differences.

As noted above, creating broader groups in terms of gender-specific differences in socio-demographic factors increased the differences in cognitive performance and aging. In the case of STM, the gender difference between peak scores increased from .21 SDs to .28 SDs. Notably, differences in verbal abilities appeared, with women reaching a peak age significantly earlier, and men having a significantly higher peak score by 0.05 of one standard deviation. However, although the gender gap was smaller (or absent) in the matched sample, this does not mean that differences in the unmatched sample should be ignored. While they may not necessarily be inherent to biology, environmental influences are a part of life, and they do drive gender differences in cognitive abilities. Thus, it is reasonable to conclude that gender differences in cognition, based on biological sex alone, are minimal; however, there are notable effects of environmental factors that in turn drive gender differences in cognition.

One large area of disparity that remained even when controlling for environmental factors was with respect to the age at which reasoning abilities began to decline. Women declined significantly earlier than men, even when controlling for demographic factors. We were also not able to capture a reliable measure of the age at which reasoning abilities peak in either gender. In women, scores declined from 12 years of age. This could be because 12 is the age at which women’s reasoning abilities do indeed peak. However, it is also possible that women peak earlier, but due to lack of data we were unable to determine the true peak from the current sample. Similarly, both unmatched and matched samples of men showed a plateau in reasoning scores until the point at which they began to decline. There are several possible explanations here. First, it is possible that men do peak in early adulthood, somewhere between 18 and 24 years of age, but the increase in reasoning abilities was not captured due to too small a sample size or noisy data. Second, they could follow a similar trajectory to women, with a slow decline before a steeper one, again not captured due to a lack of data. Because our sample of men was very large (over 32,000 in the unmatched sample), it is unlikely that either of these options are the case. Third, this plateau could be a true peak in reasoning, lasting several years, before beginning to decline. Previous research does suggest that reasoning abilities are relatively mature by age 12 (Anderson, 2002; Anderson et al., 2001b), and another large-scale study has shown that by age 18, reasoning abilities have begun to decline (Salthouse, 2009). Thus, although it is not possible to confirm that decline begins around age 12 in the current sample of women, the data follow a pattern that fits previous research and supports this claim.

The results presented here offer some insight into how to tailor interventions for cognitive decline appropriately for each gender. For example, women are known to experience more anxiety than men (McLean et al., 2011), a fact reflected in the current sample. Anxiety is known to correlate negatively with working memory (Moran, 2016). Thus, to improve working memory, or protect against its decline, therapies should perhaps focus on reducing anxiety in everyone, with a targeted focus on women. Another example is substance abuse, which is more prevalent in men (Compton et al., 2007). Because substance abuse negatively affects cognition (Crego et al., 2009), especially with respect to aging (Woods et al., 2016), a focused campaign aimed to reduce drug and alcohol consumption in men may yield a slowing in cognitive decline at the male population level. These gender-focused interventions can be combined with other treatments known to provide protection from cognitive decline, such as frequent exercise (Erickson et al., 2011) for a well-rounded defence against cognitive aging.

## Acknowledgements

This research was funded by a Canada Excellence Research Chair (CERC) program grant (#215063) to A.M.O., an AGE-WELL NCE and Women’s Brain Health Initiative (WBHI) grant to E.S.N., a Mitacs Elevate postdoctoral fellowship to E.S.N., and NSERC Discovery Grant to A.S.. A.M.O. is a CIFAR fellow.

## Declaration of Interest Statement

The cognitive tests used in this study are marketed by Cambridge Brain Sciences Inc., of which Dr. Owen is the Chief Scientific Officer. Under the terms of the existing licensing agreement, Dr. Owen and his collaborators are free to use the platform at no cost for their scientific studies and such research projects neither contribute to, nor are influenced by, the activities of the company. As such, there is no overlap between the current study and the activities of Cambridge Brain Sciences Inc., nor was there any cost to the authors, funding bodies or participants who were involved in the study.

