## Supplemental materials for "Cognition across the lifespan: age, gender, and sociodemographic influences"

**Sociodemographic, lifestyle, and sleep questionnaire**

1. What is your age?
2. Are you male or female?
3. What is your education level?
4. What is your employment status?
5. How often do you exercise (e.g., running, going to the gym)?
6. Approximately how many hours did you sleep last night?
7. How many units of alcohol do you drink each week?
8. How many caffeine containing drinks do you consume each day?
9. How many cigarettes do you smoke each day?

In your opinion, how often do you suffer from the following problems?

10. Difficulty sleeping
11. Feelings of anxiety
12. Do you consider yourself to be technologically savvy or 'out of touch with the latest gadgets'?
13. How often do you play computer games?
14. Do you consider yourself to be liberal, conservative or in the middle?
15. How religious are you?
16. How old are you relative to your siblings?

### **Test Descriptions**

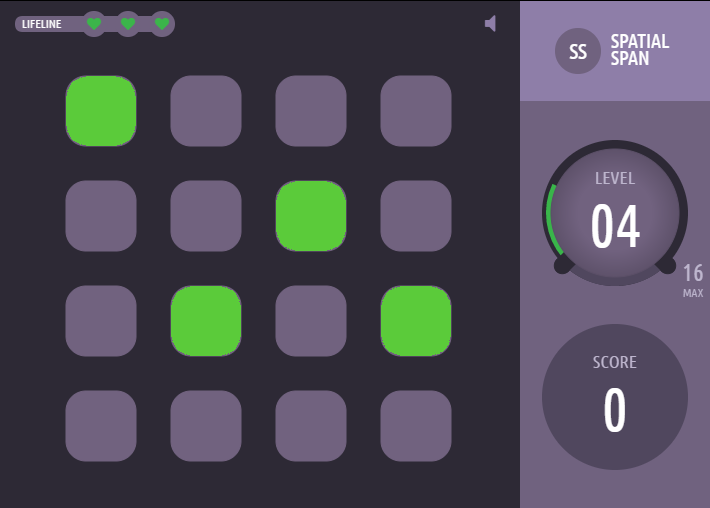

**Spatial Span** is based on the Corsi Block Tapping Task - a tool for measuring spatial short-term memory capacity. 16 purple boxes are displayed in a grid. A sequence of randomly selected boxes turn green one at a time (900 ms per green square). Participants must then repeat the sequence by clicking boxes in the same order. Difficulty is varied dynamically: correct responses increase the length of the next sequence by one square, and an incorrect response decreases the sequence length. The test finishes after 3 errors. The score is the length of the longest sequence successfully remembered.

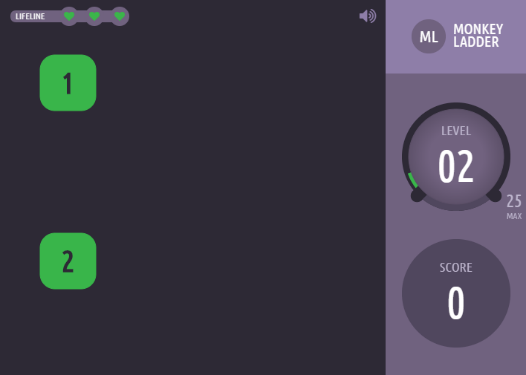
**Monkey Ladder** is based on a task from the non-human primate literature (Inoue and Matsuzawa, 2007). Numbered boxes are displayed (at the same time) at random locations within a grid. After a variable interval (number of squares * 900 ms), the numbers disappear leaving only the boxes. Participants must click the boxes in ascending numerical sequence. Difficulty is varied dynamically like in Spatial Span. The test finishes after 3 errors, and the resulting score is the length of the longest sequence successfully remembered.

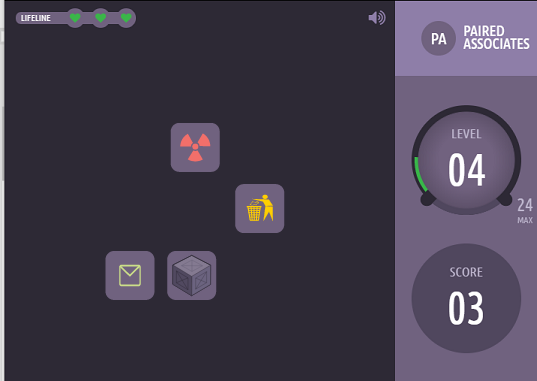
**Paired Associates** is based on a test commonly used to assess memory impairments in aging clinical populations (Gould et al., 2005). Sets of boxes are displayed at random locations on grid. The boxes open one after another to reveal an icon, after which they close. The icons are then displayed sequentially in the centre of the screen, and the participant must select box that contained that icon. If the participant remembers all the icon-location pairs correctly, then the next trial will have one more box. If an error is made the next trial has one less box. The test ends after three errors. The participant’s score is the maximum number of pairs successfully remembered.

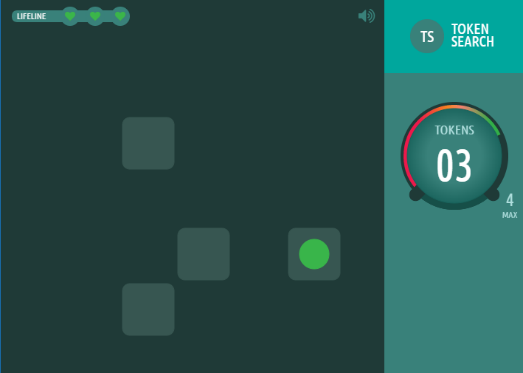
**Token Search** is based on a test that is widely used to measure strategy during search behaviour (Collins et al., 1998). A set of boxes, one of which contains a hidden green token, is displayed on a grid. Participants must find the token by clicking the boxes one at a time. Once found, it is hidden within another box. The token will not appear within the same box twice, thus, the participant must search the boxes until the token has been found once within each box. An error is made if the participant checks a box that has: 1) already been clicked while trying to find the token, or 2) previously contained the token. If the participant makes an error, a new trial begins with one less box to search. If the token is found once in each box without any errors being made, a new trial begins with one more box to search. The test finishes after three errors. The resulting score is the maximum level completed.

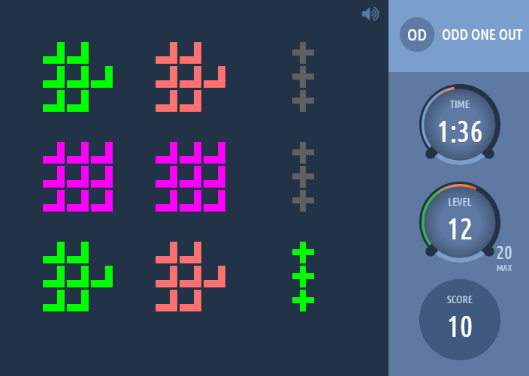
**Odd One Out** is based on a sub-set of problems from the Cattell Culture Fair Intelligence Test (Cattell, 1949). Nine groups of coloured shapes are displayed in a grid. The features define each group (colour, shape, # of items) are related to each other according to a set of rules. Participants must deduce the rules that relate these features and select the group whose contents do not correspond to those rules. They have 90 seconds to solve as many problems as possible, and the puzzles get progressively more difficult. A correct response increases the final score by one point, whereas an incorrect response decreases the score by one point.

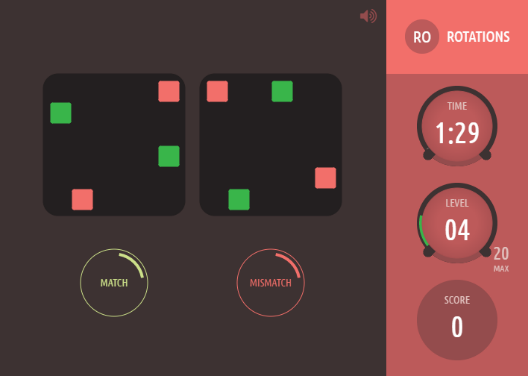
**Rotations** is a task that measures the ability to spatially manipulate objects in mind (Silverman et al., 2000). One each trial, two groups of coloured squares (each with N squares) are displayed beside each other. One of the groups is rotated by a multiple of 90 degrees. The groups are either identical (when un-rotated) or differ by the position of just one item, and participants must indicate if the groups match. They have 90 seconds to complete as many trials as possible. A correct response increases the final score by N, and the subsequent trial has groups of N+1 squares. If the response is incorrect, the total score decreases by N, and next trial has groups of N-1 squares.

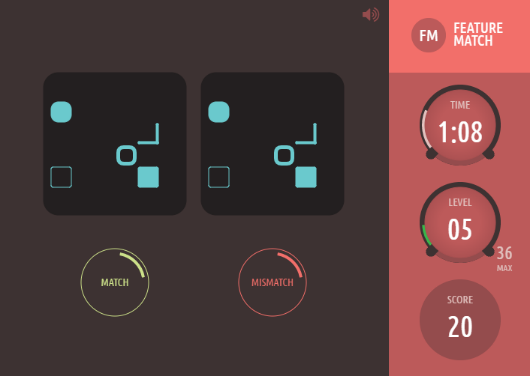
**Feature Match** is based on classic feature search tasks used to measure attentional processing (Treisman and Gelade, 1980). One each trial, two groups of items (each with N items) are displayed beside each other. The groups are either identical in their contents (and item positions), or differ by just one item. Participants have 90 seconds to complete as many trials as possible, indicating whether the groups match. A correct response increases the final score by N, and the subsequent trial has groups of N+1 items. If the response is incorrect, the total score decreases by N, and next trial has groups of N-1 items.

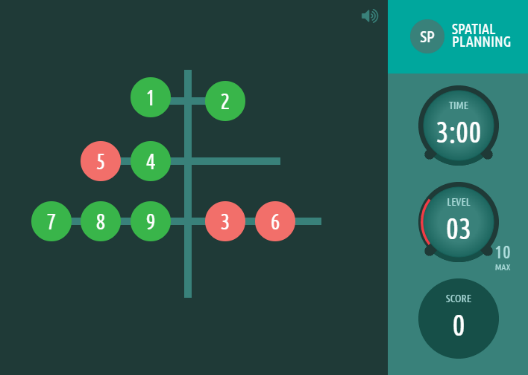
**Spatial Planning** is based on the Tower of London Task (Shallice, 1982), which is widely used to measure executive function. Numbered beads are positioned on a tree, and the participant must relocate the beads so that they are arranged in ascending numerical order. They have 3 minutes to solve as many puzzles as possible, which become progressively harder - requiring more moves and more complex planning. Trials are aborted if the participant makes more than twice the number of moves required to solve the problem. A successfully completed puzzle increases the final score by: (2 x minimum # of moves required) minus the # of moves made.

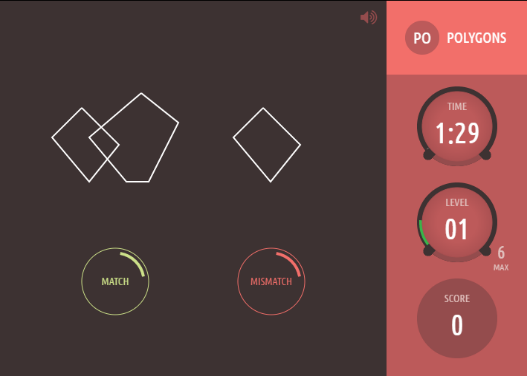
**Polygons** is based on the Interlocking Pentagons task, often used for assessing age-related disorders (Folstein et al., 1975). On each trial, two overlapping wire-framed polygons are displayed on the left side of screen, and participants must indicate whether the shape to the right is identical to one of the two overlapping ones. A correct response increases the total score by the difficulty level, and the subsequent trial will be more difficult (i.e., differences between polygons will be subtler). An incorrect response decreases the total score by the difficulty level, and the next trial will be slightly easier.

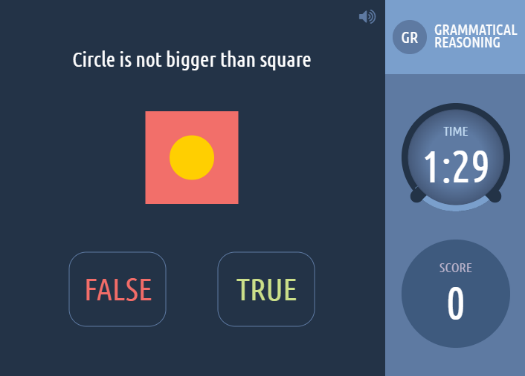
**Grammatical Reasoning** is based on Alan Baddeley’s 3-minute grammatical reasoning test (Baddeley, 1968). On each trial, a written statement regarding two shapes is displayed on the screen, and the participant must indicate whether it correctly describes the shapes pictured below. The participant has 90 seconds to complete as many trials as possible. A correct response increases the total score by one point, and an incorrect response decrease the score by one point.

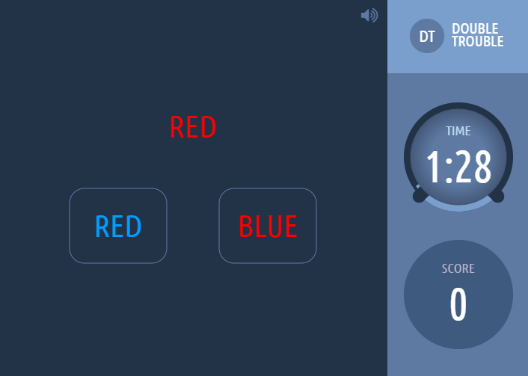
**Double Trouble** is a novel and challenging variant of the Stroop test (Stroop, 1935). A target word (either “RED” or “BLUE”) is displayed on the screen in either the colour red or the colour blue. The participant must select the probe word that correctly describes the colour that the target word is drawn in. The problem’s colour mappings can be: congruent (if every word correctly describes the colour it is displayed in); incongruent (if either the target word, or both probe words, are displayed in the opposite colour); or doubly incongruent (if the target and probes are written in the colours opposite to what they describe). Participants have 90 seconds to complete as many trials as possible. A correct response increases the total score by one point, and an incorrect response decrease the score by one point.

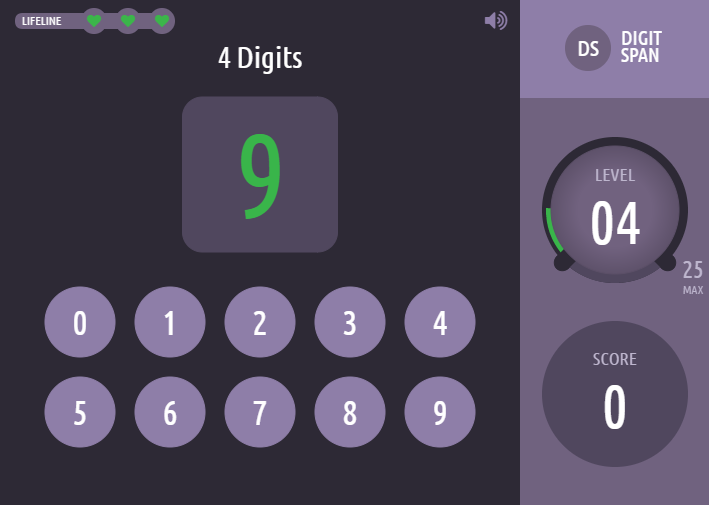
**Digit Span** is based on the verbal working memory component of the WAIS-R intelligence test (Weschler, 1981). A sequence of digits is displayed, one at a time, in green in the centre of the screen. Participants must then repeat the sequence of digits by selecting them on the on-screen keyboard. Difficulty is dynamically varied like previous tests, and the test ends after 3 mistakes. The resulting score is the length of the longest digit sequence successfully remembered.

**Supplemental Figures**

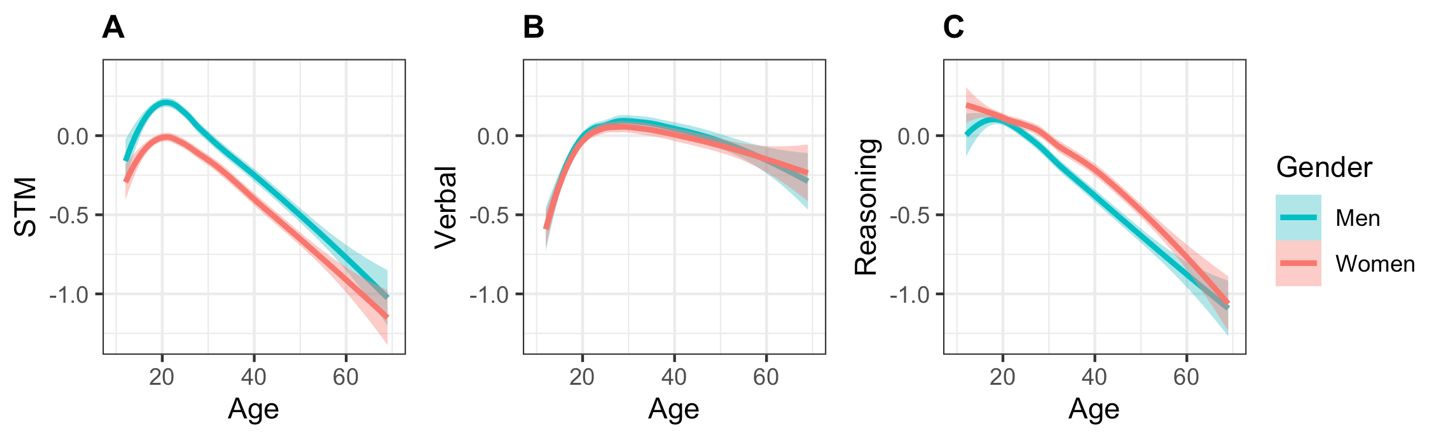
 **Fig S1.** Local regression curves for STM, Verbal, and Reasoning scores across the lifespan, ranging from 12 to 69 years of age, in N = 18,902. 95% confidence bands are shown in translucent colour around the line.

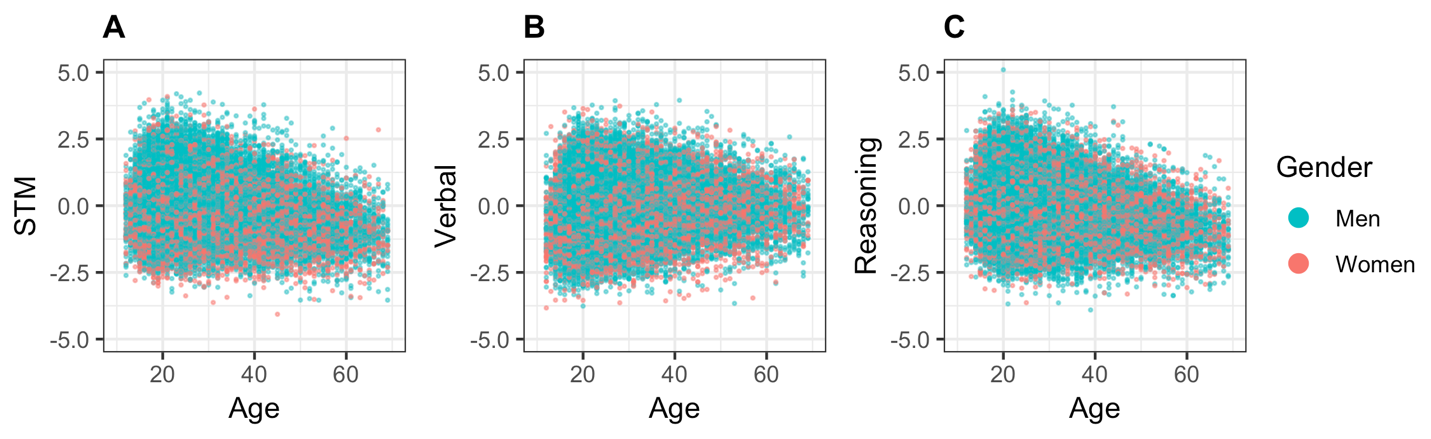

**Fig S2.** All 45,779 raw data points across age for STM (A), Verbal (B), and Reasoning (C) cognitive domains.

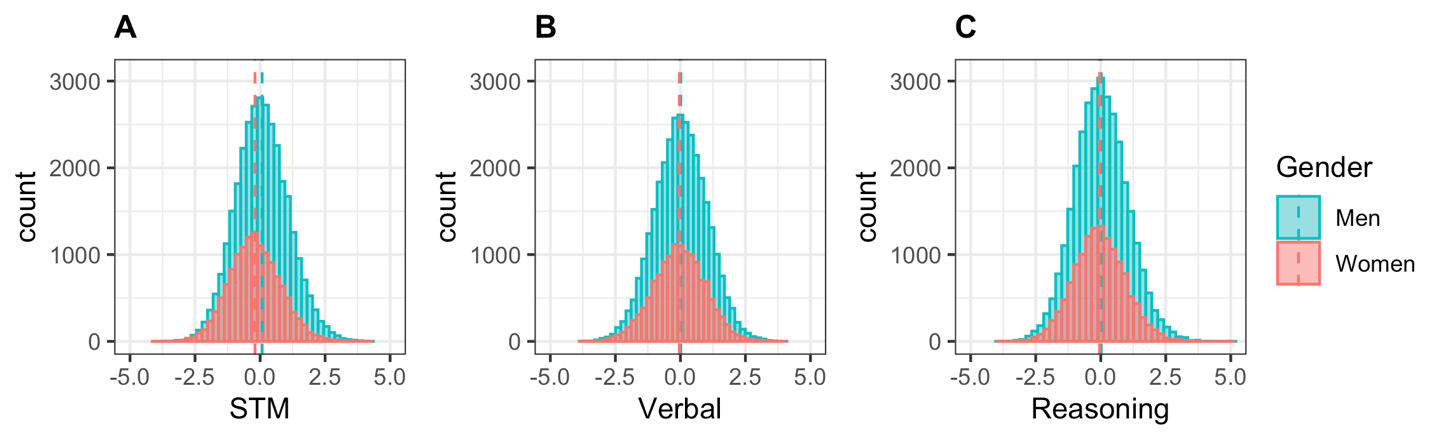

**Fig S3.** Histograms of cognitive domain scores by gender, in N = 45,779. Dashed lines indicate mean.

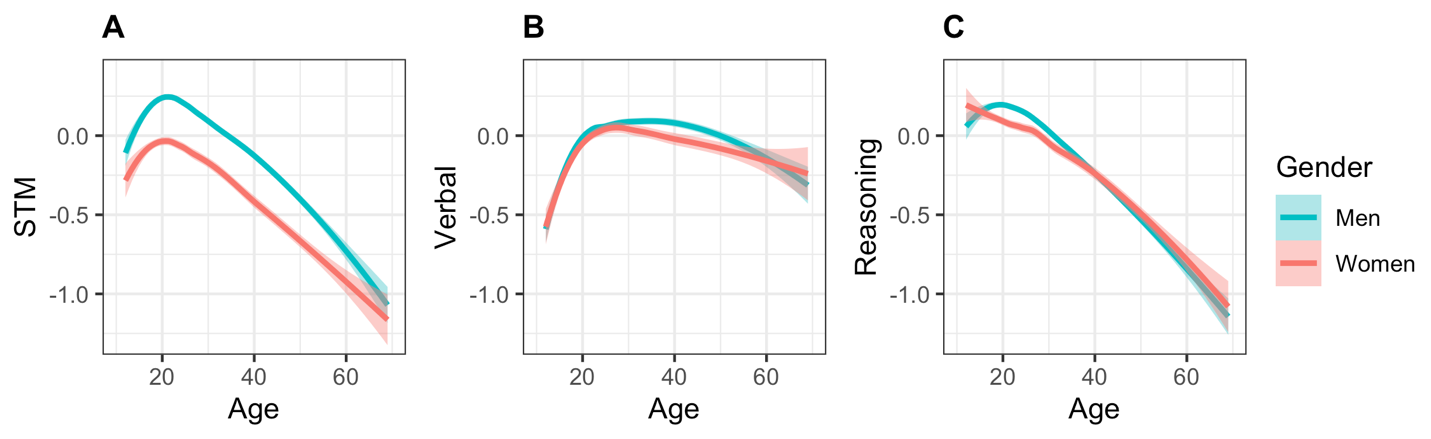

**Fig S4.** Local regression curves for STM, Verbal, and Reasoning scores across the lifespan, ranging from 12 to 69 years of age, in N = 45,779. 95% simultaneous confidence bands are shown in translucent colour around the line.

**Supplemental Tables**

| **Table S1**. **Comparison of demographic variables across women and men.** | | | | | | |
| --- | --- | --- | --- | --- | --- | --- |
| Measure | | Mean (SD) or Percentage | | *t*(df), W, χ^2^(df, N)^1^ | *p* | Effect size^2^ |
|  |  | Women | Men |  |  |  |
| *N* | | 13,444 | 32,335 |  |  |  |
| Age (years) | | 28.08 (11.01) | 28.22 (10.30) | -1.31(23,696) | 0.220 | -0.01 |
| Sleep (hours per night) | | 7.09 (1.64) | 6.93 (1.64) | 9.51(25,241) | < .001 | 0.10 |
| Alcohol (units per week) | | 1.72 (1.85) | 1.78 (2.00) | -3.10 (27,022) | .003 | -0.03 |
| Caffeine (units per day) | | 3.24 (4.69) | 4.22 (5.53) | -19.36 (29,414) | < .001 | -0.19 |
| Cigarettes (per day) | | 1.48 (4.54) | 1.71 (5.06) | -4.68 (27,830) | < .001 | -0.05 |
| Highest education completed | |  |  | 2.17e^8^ | .989 | 6.27e^-5^ |
|  | Some high school | 10.20% | 9.80% |  |  |  |
|  | High School | 8.40% | 10.60% |  |  |  |
|  | Some post-secondary | 27.80% | 30.50% |  |  |  |
|  | Post-secondary degree | 27.20% | 27.70% |  |  |  |
|  | Professional degree | 26.40% | 21.40% |  |  |  |
| Level of employment | |  |  | 2.21e^8^ | .003 | 0.01 |
|  | No answer | 4.10% | 2.70% |  |  |  |
|  | Unemployed | 10.30% | 12.10% |  |  |  |
|  | Full time student | 28.70% | 25.40% |  |  |  |
|  | Employed and student | 15.50% | 11.90% |  |  |  |
|  | Employed part time | 10.00% | 6.20% |  |  |  |
|  | Employed full time | 31.40% | 41.80% |  |  |  |
| Exercise | |  |  | 2.18e^8^ | .974 | 5.35e^4^ |
|  | Never | 10.60% | 11.00% |  |  |  |
|  | Infrequently | 37.80% | 34.70% |  |  |  |
|  | Weekly | 19.90% | 18.20% |  |  |  |
|  | Several times a week | 25.30% | 28.30% |  |  |  |
|  | Every day | 6.50% | 7.90% |  |  |  |
| Depressive feelings | |  |  | 2.36e^8^ | < .001 | 0.07 |
|  | No answer | 1.10% | 1.10% |  |  |  |
|  | Never | 10.40% | 17.50% |  |  |  |
|  | Occasionally | 56.80% | 54.00% |  |  |  |
|  | Quite often | 21.40% | 18.20% |  |  |  |
|  | Nearly every day | 7.30% | 6.40% |  |  |  |
|  | All the time | 3.10% | 2.80% |  |  |  |
| Anxiety | |  |  | 2.58e^8^ | < .001 | 0.15 |
|  | No answer | 1.30% | 1.50% |  |  |  |
|  | Never | 13.00% | 24.00% |  |  |  |
|  | Occasionally | 48.80% | 49.90% |  |  |  |
|  | Quite often | 21.20% | 14.80% |  |  |  |
|  | Nearly every day | 10.80% | 6.50% |  |  |  |
|  | All the time | 5.00% | 3.20% |  |  |  |
| Video games | |  |  | 2.42e^9^ | < .001 | 0.09 |
|  | Never | 38.00% | 17.10% |  |  |  |
|  | Monthly | 26.00% | 20.30% |  |  |  |
|  | Weekly | 21.40% | 29.40% |  |  |  |
|  | Daily | 14.60% | 33.20% |  |  |  |
| Siblings | |  |  | 2.19e^8^ | .111 | 0.01 |
|  | Only child | 12.40% | 10.60% |  |  |  |
|  | Youngest | 30.40% | 31.00% |  |  |  |
|  | Middle | 16.40% | 17.40% |  |  |  |
|  | Oldest | 40.80% | 41.00% |  |  |  |
| Religiosity | |  |  | 2.34e^8^ | < .001 | 0.06 |
|  | Atheist | 31.90% | 42.40% |  |  |  |
|  | Agnostic | 31.50% | 31.50% |  |  |  |
|  | Religious lapsed | 19.50% | 15.10% |  |  |  |
|  | Religious practicing | 13.10% | 8.10% |  |  |  |
|  | Very religious | 3.90% | 2.90% |  |  |  |
| Political leaning | |  |  | 68.67(2,  *N* = 45,779) | < .001 | 0.04 |
|  | Liberal | 47.20% | 44.90% |  |  |  |
|  | Middle | 45.20% | 45.10% |  |  |  |
|  | Conservative | 7.60% | 10.00% |  |  |  |
| Tech savvy | |  |  | 2693.40(1, *N* = 45,779) | < .001 | 0.24 |
|  | Yes | 64.00% | 89.10% |  |  |  |
|  | No | 36.00% | 10.90% |  |  |  |
| ^1^Welch’s *t*-test used to compare numeric variables, Wilcoxon Rank-Sum used to compare ordinal variables, and χ^2^  used to compare categorical variables | | | | | | |
| ^2^Effect sizes used were Cohen’s *d* for t-tests, *r* for Wilcoxon Rank-Sum tests, and Cramer’s *V* for χ^2^  tests | | | | | | |
